# PROTECTIVE EFFECTS OF TRITERPENOID BETULIN ON TYPE 2 DIABETES MELLITUS IN RATS

**DOI:** 10.1101/2023.07.27.550802

**Authors:** A. H. Shlyahtun, Yu. Z. Maksimchik, A. Zakrzeska, I. P. Sutsko, A. F. Raduta, E. V. Buksha, E. V. Bogdevich, P. Kitlas, M. Tomulewicz

## Abstract

Type 2 diabetes mellitus (T2DM) is a complex chronic metabolic disease characterized by long-term hyperglycemia, which is, in turn, resulted from the impaired insulin signaling caused by a combination of insulin resistance or inadequate insulin production. Prevalence and incidence of T2DM are increasing dramatically across the world, and it is accompanied with severe complications and premature mortality of patients with diabetes. Given the fact that synthetic drugs have disadvantages in view of the side effects, the implementation of naturally occurring compounds for diabetes treatment may be a promising alternative. Betulin is a naturally occurring triterpenoid which has been shown to possess the ability of altering body lipids and exert hypoglycemic and hepatoprotective effects. It is suggested that the application of betulin in T2DM may have a favorable effect to ease the severity of diabetic complications. Thus, the aim of the present study was to assess biological effects of betulin in T2DM conditions.

Diabetes-induced rats were administered with two different doses of betulin for 28 consecutive days. It was shown that long-term administration of betulin at the doses of 50 and 100 mg/kg/day to the rats prevented diabetes-associated changes in a body weight of the animals, significantly reversed insulin resistance and abolished the impairment of glucose metabolism. It was accompanied with the dose-dependent normalization of serum lipid contents. Histopathological changes and structural abnormalities in the liver of diabetics were restored by the administration of betulin. Also, betulin was able to restrain systemic inflammation detected in diabetic animals according to the altered levels of serum TNFα. Thus, the results obtained in the current study were found to be in agreement with earlier findings on beneficial effects of betulin in conditions pathogenetically close to T2DM. We hypothesized that the ability of betulin to restrain systemic inflammation and to normalize the lipid metabolism can explain improved insulin resistance and glycemic control and it can provide a possible mechanism for the beneficial antidiabetic effects of betulin.

## Introduction

According to current knowledge, type 2 diabetes mellitus (T2DM) is described as a complex metabolic condition that is characterized by long-term hyperglycemia followed by insulin resistance and relative lack of insulin production. It is emphasized that T2DM can cause severe complications, either of microvascular (retinopathy, nephropathy and neuropathy) or macrovascular (coronary heart disease, cardiomyopathy, arrhythmias and sudden death) origin.

There has been a huge body of evidence that the spread of T2DM has become pandemic over the last few decades. Lifestyle changes as well as low-levels of physical activity and nutrient imbalances are often mentioned as the main reasons leading to the wide distribution of diabetic pathology. The World Health Organization has recently reported that by the year of 2025 there will be about 250 million people all over the world suffering with diabetes type 2. It is worth mentioning that among the main types of diabetes T2DM already accounts for about 90 % of all observations.

There are lots of obstacles that are encountered in attempting to manage diabetic pathology. At the early stages of the disease, a strict diet and physical exercises are usually prescribed. If adequate blood sugar is not achieved pharmacological correction follows-up to lower blood glucose levels. The list of oral hypoglycemic drugs used to retain blood glucose is extensive enough and includes metformin, sulfonylurea derivatives, dipeptidylpeptidase-4 inhibitors, glucagon-like peptide-1 receptor agonists, sodium-glucose cotransporter-2 inhibitors, glucosidase inhibitors and pioglitazone. If none of these pharmaceuticals were effective for controlling blood sugar, insulin injections would be used.

Despite the progress that was made in the treatment of T2DM, patients mostly are rare to meet the required glycemic control. Furthermore, contraindications to using some drugs could be the reason to change the usual treatment regimen for patients with high glucose levels and it is owing to the fact that the safety and efficacy of some novel hypoglycemic agents are still under question. On the other hand, severe disorders associated with heart and respiratory failure, the occurrences of ketoacidosis or lactic acidosis, as well as conditions due to pregnancy and lactation could be the reason to postpone the treatment. Ultimately, it is well known that the side effects of commonly used hypoglycemic drugs may affect digestive system (nausea, vomiting, diarrhea), endocrine system (hypoglycemia) and provoke metabolic changes (acidosis) [1].

Taking into account the struggles of diabetes management discussed earlier, it is obvious that an effective strategy for prevention and treatment of T2DM is still required. Given the fact that synthetic drugs have disadvantages in view of the side effects, the implementation of naturally occurring compounds for diabetes treatment may be a promising alternative.

Betulin is natural pentacyclic triterpenoid (lup-20(29)-ene-3β,28-diol) (Figure 1) found out in nature in several plants. There is a growing interest in betulin accounted for its wide spectrum of biological activities. Betulin is usually extracted from the outer bark of the birch tree and the calculations show that this substance represents about 40 % of dry matter. Pharmacological properties of betulin are being intensively studied *in vivo* and *in vitro*. It has been shown that this substance can exert antitumor, antimicrobial, antioxidant, anti-inflammatory, immunomodulatory, and hepatoprotective effects [2].

**Figure 1.**
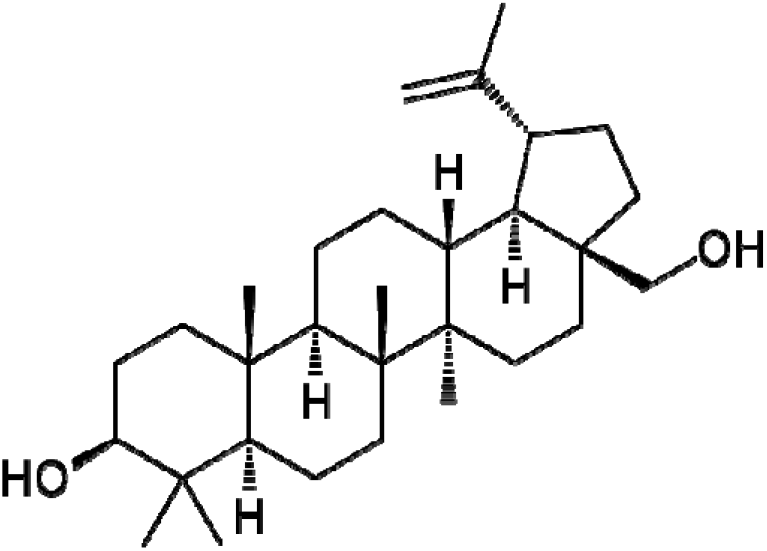
The chemical structure of betulin.

Recently, we demonstrated that betulin was effective to normalize lipid metabolism and alleviate oxidative stress in pathological conditions close to T2DM including alcoholic and non-alcoholic fatty liver disease [3]. Owing to the fact that development of T2DM is usually associated with obesity and impaired lipid turnover, the alleviation of diabetic dyslipidemia is considered to be the important part of therapeutic intervention. It is suggested that the ability of betulin to affect the intensity of lipid exchange may have a favorable effect to ease the severity of complications in T2DM.

Thus, the aim of the present study was to evaluate the effects of betulin on glycemic levels, lipid contents, and histology of liver in rats with experimental T2DM and to elucidate the specific mechanisms by which betulin may improve diabetes management.

## Materials and methods

Unless otherwise stated, all chemicals used in the experiment were of reagent-grade quality. Analytical standard of betulin was obtained from Carl Roth (No. 8763.1, Germany). Streptozotocin was from Sigma-Aldrich (S0130, USA). Organic solvents were of analytical grade and used without further purification. Merck Millipore Direct Q3 system (USA) was used to produce deionized water to prepare buffer solutions.

Outer bark of birch (*Betula pendula Roth*.) was collected in the forest located around Grodno, Belarus. The origin of the plant materials was confirmed by qualified botanist at Grodno State University, Belarus.

Betulin was isolated from the birch bark by Soxhlet extraction following on the method described earlier [4]. Attenuated total reflectance Fourier-transform infrared spectroscopy technique was recru ted for qualitative analysis of betulin in extracts. The samples were analyzed with the Thermofisher Scientific Nicolet iS 10 IR Fourier spectrometer (USA) with the measurements made in the main absorption region (4000 to 400 cm^-1^). The purity of betulin obtained during the isolation was 98,9 % as evidenced reverse-phase chromatography analysis on the Agilent 1200 system using the method described by Zhao et al. [5].

The animal study was carried out on the 80 male Wistar rats. The animals were kept in the plastic cages containing 8 rats in the cage and had free access to food and water. The temperature, humidity, and a light cycle were controlled in the room where the animals were housed.

In order to develop T2DM in rats, slightly modified standard HFD/STZ protocol was used [6]. According to the protocol, the rats were provided with the high fat diet (HFD) for 42 days. The main component of the diet was a mixture of animal fats covering 45 % of nutritional values in nourishment to be provided to the rats. Additionally, to ensure diabetic conditions the animals were injected with streptozotocin two times on the 14^th^ and 28^th^ day since HFD had been started. Streptozotocin was dissolved in a 100 mM sodium-citrate buffer with a pH of 4.4 and injected at a dose of 25 mg/kg. Blood glucose levels (pre-prandial glucose) was measured following the 42 days period of HFD and the animals with glucose more than 12 mmol/l were used for further experimental procedures.

To proceed the experiment, the rats were divided into 5 groups which were labeled as “Control” (12 rats), “Betulin” (12 rats), “T2DM” (12 rats), “T2DM + Betulin 50” (10 rats), “T2DM + Betulin 100” (10 rats).

The animals included in “Control” and “Betulin” groups were kept on the standard diet during the whole time of the experiment duration. Similarly, the rats that belonged to “T2DM”, “T2DM + Betulin 50” and “T2DM + Betulin 100” groups received HFD during the whole duration of the experiment.

The rats included in “T2DM + betulin 50” and “T2DM + betulin 100” groups received betulin intragastrically at the doses of 50 and 100 mg/kg/day, respectively, for consecutive 28 days. Owing to low water solubility, a suspension of betulin in 2 % gelatinized starch was constantly prepared in order to facilitate the administration of the substance to the animals.

During the whole experiment, pre-prandial blood glucose levels were weekly monitored with Roche Diabetes Care Accu-Chek Active Blood Glucometer (Germany).

The oral glucose tolerance test (OGTT) followed by the evaluation of glucose area under curve (AUC) was done twice as an index of glucose intolerance. First time this test was performed after the formation of the experimental groups. Second time it was carried out at the end of the experiment to assess the effect of betulin on the development of insulin resistance. To do the test and further calculations, the animals were administered intragastrically with a 40 % glucose solution at a dose of 2 g/kg. Blood glucose levels were assayed immediately after the glucose administration (0 min), 30, 60, and 120 min since the beginning of the test. Based on the glucose measurements AUC was calculated for each of the animals. Calculations were done following the method of trapezoidal approximation of postprandial blood glucose levels with the aid of GraphPad Prism v. 8.0 software (USA) [7].

At the end of the experiment, the rats were euthanized and blood and tissue samples were collected. Blood samples were centrifuged at 2000 g for 10 min to obtain blood serum which was stored in a freezer at −82 °C until analysis. Liver tissues to take into histological study were fixed with 10 % formalin. Samples thus obtained were dehydrated by means of immersing them in a series of alcohols of increasing concentration and embedded in paraffin wax. Following that, histological sections were prepared and on dewaxing them stained with hematoxylin and eosin. For further microscopic analysis, liver sections were imaged with the microscope Leica DM6 B (Germany). Photomicrographs thus prepared were processed with computer software. The intensity of the inflammation process in liver tissues was assessed counting the number of foci of the lymphocytic infiltration. The intensity of inflammatory processes in the liver was assessed by the number of foci of lymphocytic infiltration which is defined as three or more lymphocytes detected in ten fields of view of the microscope.

Serum activities of alanine aminotransferase (ALT) and aspartate aminotransferase (AST) as well as the levels of total cholesterol (TC), triacylglycerols (TG), and high-density lipoprotein cholesterol (HDL-C) were estimated with commercially available kits from AnalysX (Belarus). Enzyme-linked immunosorbent assay kits for insulin, glycated hemoglobin, and tumor necrosis factor α (TNFα) measurements were purchased from Cusabio (China). Free fatty acids (FFA) contents in serum of rats were obtained following the method of Duncombe [8].

The European Convention for the Protection of Vertebrate Animals used for Experimental and other Scientific Purposes was followed on each step of the study (ETS No.123).

Statistical analysis of the data obtained was carried out with the aid of GraphPad Prism v. 8.0 software (USA). The Shapiro-Wilk test was used to examine whether the data sets were fitted to a normal distribution. To analyze the intragroup dynamics for the metabolic processes being studied, the statistical significance of the changes was evaluated by means of comparing the average values of the test results obtained at the beginning and at the end of the study using two-tailed Student’s t-test. To identify the differences between the experimental groups, the analysis of variance and the post-hoc Tukey test were used. All results were expressed as a mean value followed by standard error (M ± m).

## Results and discussion

Feeding the rats with HFD for 42 days and two injections of STZ resulted in significant changes in body weight, fasting blood glucose and AUC when compared to those without STZ injection and kept on a standard diet. It was noticed that HFD and STZ increased body weight of the rats by 17.7 % (p<0.05) including those animals that would later obtain different doses of betulin. We also found the elevation in fasting blood glucose levels (the average value was 12.1±1.7 mmol/l) and impaired glucose tolerance according to results of OGTT (a 53.25 % increase in glucose AUC was observed which corresponded the changes up to 26.7±5.1 mmol×h/l). To sum up, the data obtained approved that the experimental animals had developed T2DM before administration of betulin was started (Table 1).

**Table 1.**
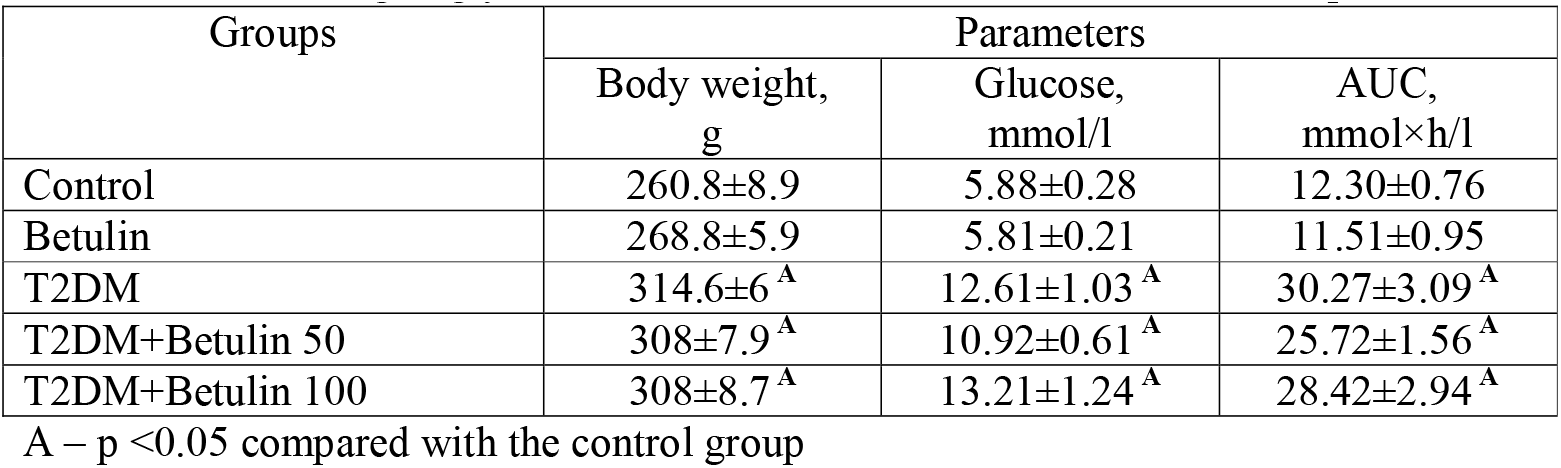
Animal weight, glycemic levels and OGTT results at the start of experiment.

| Groups | Parameters |  |  |
| --- | --- | --- | --- |
| | Body weight, g | Glucose, mmol/l | AUC, mmol×h/l |
| Control | $260.8 \pm 8.9$ | $5.88 \pm 0.28$ | $12.30 \pm 0.76$ |
| Betulin | $268.8 \pm 5.9$ | $5.81 \pm 0.21$ | $11.51 \pm 0.95$ |
| T2DM | $314.6 \pm 6^A$ | $12.61 \pm 1.03^A$ | $30.27 \pm 3.09^A$ |
| T2DM+Betulin 50 | $308 \pm 7.9^A$ | $10.92 \pm 0.61^A$ | $25.72 \pm 1.56^A$ |
| T2DM+Betulin 100 | $308 \pm 8.7^A$ | $13.21 \pm 1.24^A$ | $28.42 \pm 2.94^A$ |

A – $p < 0.05$ compared with the control group

In the next stage of the study, the control and diabetic rats were divided and additional experimental groups were formed which followed by the administration of two doses of betulin for 28 consecutive days. At the end of the experiment before the rats were sacrificed their body weight and blood glucose levels were measured, and OGTT was performed.

As a result, the animals in T2DM group had body weight significantly higher (by 20.5 %) compared to controls. Blood glucose was also significantly increased in diabetics reaching the value of 13.4±1.1 mmol/l (an increase by 124.5 %). Diabetic condition of experimental animals was also confirmed by estimating blood levels of glycated hemoglobin which appeared to be by 71.9 % [up to 9.8±0.4 % (p <0.001)] higher in diabetic animals compared to control group. The increase in formation of glycated hemoglobin levels perfectly described hyperglycemic condition of diabetic animals and indicated on increased rates of formation of glycated proteins by non-enzymatic reactions.

In addition to hyperglycemia, diabetic rats had developed insulin resistance as evidenced by 2.6 times increase in AUCs of glucose tolerance test and 2.8 times increase of serum insulin levels in comparison to control rats (Table 2).

**Table 2.** Effect of betulin administration on body weight, glycemic levels, OGTT results, glycated hemoglobin, and insulin levels in the blood serum of rats with T2DM.

| Groups | Body weight, g | Glucose, mmol/l | AUC, mmol×h/l | Hb1A, % | Insulin, pg/ml |
| --- | --- | --- | --- | --- | --- |
| Control | $295.4 \pm 10.6$ | $5.96 \pm 0.23$ | $12.28 \pm 0.79$ | $5.7 \pm 0.29$ | $50.5 \pm 2.2$ |
| Betulin | $291.3 \pm 5.8$ | $5.76 \pm 0.22$ | $11.51 \pm 0.95$ | $5.8 \pm 0.26^B$ | $69.1 \pm 3.7$ |
| T2DM | $355.8 \pm 9.9^A$ | $13.39 \pm 1.08$ | $31.44 \pm 4.47^A$ | $9.8 \pm 0.43^A$ | $141.0 \pm 22.3^A$ |
| T2DM+Betulin 50 | $327.0 \pm 10.6^A$ | $11.02 \pm 0.76$ | $22.64 \pm 1.51^{AB}$ | $8.9 \pm 0.45^A$ | $74.0 \pm 5.2^{AB}$ |
| T2DM+Betulin 100 | $309.5 \pm 6.1^B$ | $8.06 \pm 0.40^B$ | $16.14 \pm 1.48^B$ | $7.2 \pm 0.26^{AB}$ | $62.0 \pm 3.3^B$ |

A – $p < 0.05$ compared with the control group, B – $p < 0.05$ compared with the T2DM group

The administration of betulin to diabetic rats led to a decrease in the body weight of animals and this effect was dose-dependent. The severity of hyperglycemia as well as insulin resistance estimated by glucose AUCs and serum insulin levels were also alleviated in the same manner. In particular, it was shown that blood glucose levels were lowered by 17.7 % in diabetic animals received 50 mg/kg of betulin per day, and it was 39.8 % decrease in blood glucose of those received 100 mg/kg/ of betulin daily compared to untreated T2DM group. Similarly, glucose AUC in diabetic rats administered with betulin was found to be declined by 28.0 % and 48.7 % respectively compared to untreated animals with T2DM. Finally, the most noticeable changes revealed the data on insulin levels showing that higher dose of betulin not only prevented manifestations of diabetes but reduced this marker of diabetic pathology to normal values.

It is well known that T2DM is followed by specific changes in lipid metabolism which results in severe disturbances of serum lipids and subsequent dyslipidemia [9]. Thus, the key components involved in lipid exchange were measured in blood plasma of experimental animals in order to clarify the effects of betulin on lipid status.

It was shown that the lipid levels in blood plasma of control animals and in non-diabetic rats administered with betulin were within the scopes of physiological ranges attributed to Wistar rats [10].

Rats with T2DM were found to reveal the development of severe dyslipidemia. There was an increase in TC content by 73.8 % and a decrease in HDL levels by 21.5 % compared to control animals. Additionally, TG content in the blood of rats with T2DM was 1.7 times higher than that in the control group. We also found elevated levels of FFA in the diabetic group (Table 3). At our knowledge, long-term exposition to high concentrations of FFA can produce a toxic effect usually referred as lipotoxicity. Increased serum FFA concentrations affect adipocytes, cardiomyocytes, hepatocytes, and pancreatic β-cells leading to impairments of insulin signaling and development of insulin resistance in target tissues [11].

**Table 3.** Indicators of lipid metabolism in serum of rats with T2DM administered with betulin, mmol/l.

| Groups | Parameters |  |  |  |
| --- | --- | --- | --- | --- |
|  | TC | HDL-C | TG | FFA |
| Control | 4.31±0.49 | 3.12±0.39 | 0.71±0.05 | 0.53±0.04 |
| Betulin | 5.12±0.57 | 4.27±0.50<br>p(control)=0.085 | 0.72±0.07 | 0.44±0.02<br>p(control)=0.051 |
| T2DM | 7.49±0.75 <sup>A</sup> | 2.45±0.32 | 1.90±0.11 <sup>A</sup> | 0.71±0.06 <sup>A</sup> |
| T2DM+Betulin 50 | 5.30±0.57 | 2.33±0.21 | 1.20±0.08 <sup>AB</sup> | 0.69±0.04 <sup>A</sup> |
| T2DM+Betulin 100 | 4.32±0.84 <sup>B</sup> | 2.71±0.37 | 0.89±0.14 <sup>B</sup> | 0.61±0.03 <sup>B</sup> |

A – p <0.05 compared with the control group, B – p <0.05 Compared with the T2DM group

Based on our findings we may suggest that diabetic animals in our study revealed a specific type dyslipidemia found earlier in diabetic patients and referred to as “lipid triad” [12, 13]. The term is used to describe three lipid abnormalities involving an increase in the concentrations of TG and LDL and a decrease in the levels of HDL and currently it is accepted to be a marker of T2DM in humans.

The administration of two different doses of betulin to rats with experimental T2DM was accompanied with the dose-dependent normalization of total cholesterol levels and HDL, TG and FFA contents (Table 2).

It is interesting to note that non-diabetic animals administered with betulin showed a tendency toward increment in serum HDL content (by 36.9 %, p<0.1) while the FFA content was slightly decreased (by 16.7 %, p=0.051) compared to control rats.

Liver is the key metabolic participant in living organisms and it plays an important role in lipid and carbohydrate metabolism. Needless to say, that it is responsible for the synthesis and storage of glycogen in order to maintain blood glucose levels in normal. According to clinical reports, T2DM associated liver disorders were found in 35-100 % of observations including the occurrence of non-alcoholic fatty liver disease which ranged between 30–80 % [14].

Following the idea of comprehensive assessment of the betulin influence on hepatic tissues and looking for its hepatoprotective effects, the histological examination of livers of experimental animals was performed.

The histological structure of livers of control rats was typical for this animal species without signs of pathological changes in liver parenchyma. The hepatocytes were polygonal in shape with weakly stained eosinophilic or amphiphilic granular cytoplasm and contained oval nuclei located predominately at the center of the cell. Lymphocytic infiltration was not detected in the livers of the control rats (Figure 2.1, Table 4).

**Table 4.** The number of inflammatory foci in the liver parenchyma of rats with T2DM administered with betulin.

| Groups | The number of inflammatory foci |
| --- | --- |
| Control | none |
| Betulin | none |
| T2DM | 12.5±2.1 |
| T2DM+Betulin 50 | 8.5±2.2 |
| T2DM+Betulin 100 | 5.4±1.6* |

\* – p <0.05 compared with the T2DM group

**Figure 2.**
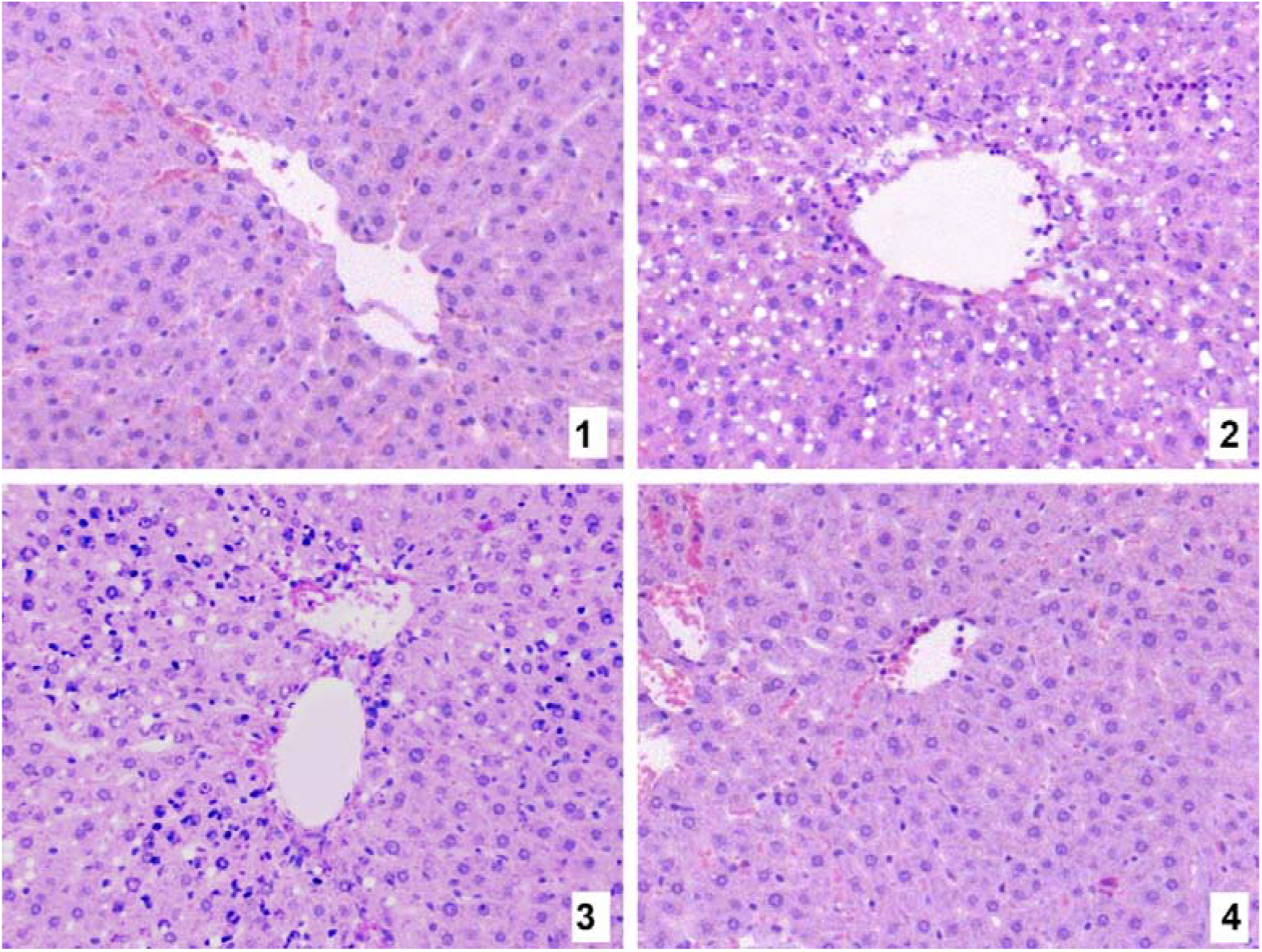
Photomicrographs of liver parenchyma of rats with T2DM Designations: 1 − Control group; 2 − T2DM; 3 − T2DM+ Betulin 50; 4 − T2DM+Betulin 100. Stained with hematoxylin & eosin. Mag. ×100.

The livers of non-diabetic animals administered with betulin seemed to have the same histological features as control rats. There were no changes in parenchyma as well as lymphocytic infiltration was no detected (Table 4).

Histological examination of the livers of HFD/STZ rats pointed out on the occurrence of microvesicular steatosis (Figures 2.2-2.4). The severity of this pathological condition varied from mild to moderate. Ballooning degeneration of hepatocytes was spotted and that was accompanied with intralobular multifocal lymphocytic infiltration. In general, the groups of 5-10 cells containing large lipid droplets were observed in the portal areas of the liver tissues. Enlarged size of the hepatic cells led the lumen of sinusoids to be narrowed. The inflammatory changes in the liver tissues resulted in lymphocytic infiltration, which was mostly seen in the areas close to the lumen of the central hepatic veins (Figure 2.2, Table 4).

The administration of betulin reduced diabetes-associated liver damage and adverted degenerative processes in the liver tissues. The higher dose of betulin was noticeably more effective to prevent pathological manifestations in the livers of diabetic rats. Both doses were able to improve diabetes-induced liver histological changes by gradually decreasing the severity of liver dystrophy and diminishing areas of ballooning degeneration in hepatic sections. The number of inflammatory foci and lymphocytic infiltration in the liver was mostly unaffected with the administration of betulin at lower dose (Figure 2.3, Table 4), while the higher dose of betulin significantly reduced inflammatory signs in liver parenchyma (Figure 2.4, Table 4).

Thus, the evidence was obtained that the administration of betulin to the rats with T2DM alleviated histopathological changes and structural abnormalities in the liver.

Serum enzymes such as alanine aminotransferase (ALT) and aspartate aminotransferase (AST) are very sensitive markers of liver injury and toxicity. In our study, the activities of ALT and AST were significantly higher in T2DM rats showing the increase by 51.1 % and 32.7 % respectively. Elevated ALT and AST were lowered with the administration of different doses of betulin to diabetic rats. Furthermore, the higher dose of betulin was more effective to return aminotransferases to control values. Therefore, based on the findings obtained for serum aminotransferases we suggest the beneficial effects of betulin on the livers of T2DM animals and this observation is in corroboration with recently discussed data on hepatoprotective effects of betulin (Table 5).

**Table 5.** Serum aminotransferases activities of rats with T2DM administered with betulin.

| Groups | ALT,<br>IU | AST,<br>IU |
| --- | --- | --- |
| Control | 60.94±4.03 | 112.20±8.09 |
| Betulin | 51.19±2.52 | 94.07±4.32 |
| T2DM | 92.09±6.84<sup>A</sup> | 148.90±12.56<sup>A</sup> |
| T2DM+Betulin 50 | 78.23±4.14 | 147.01±8.60<sup>A</sup> |
| T2DM+Betulin 100 | 69.13±3.53<sup>B</sup> | 112.30±6.14<sup>B</sup> |

A – p <0.05 compared with the control group, B – p <0.05 compared with the T2DM group

It has been described a dozen of interrelated pathogenetic mechanisms provoking the development and progression of T2DM. Among them, insulin resistance of target tissues, primarily the liver cells, occupies a central position [15]. Obesity is generally recognized as one of the main reasons for the development of insulin resistance which is subsequently followed by the disturbances in lipid metabolism. This involves further changes including dyslipidemia and adipocyte dysfunctions as well as infiltration of adipose tissues by macrophages leading to the development of systemic inflammation.

The upregulation of inflammatory reactions requires the involvement of pro-inflammatory cytokines and among them TNFα is considered to be the main component involved in the pathogenetic mechanisms of insulin resistance associated with T2DM. It is well known that elevated levels of proinflammatory cytokines make a significant contribution to the development of tissue resistance to insulin at the molecular level. Earlier it was shown that TNFα reduced the expression of glucose transporter type 4 found predominantly in adipocytes, skeletal and cardiac cells. The phosphorylation of serine residues in IRS-1 triggered by the activity of TNFα can act as an inhibitor of insulin receptor and may block signal transduction down-stream of PI3K [16].

Taking into account the information mentioned above, the measurements of serum TNFα were taken in rats with T2DM. As expected, the impact of pathological changes in rats with experimental diabetes manifested itself in a rise of TNFα production. It was revealed a significant increase (2.2 times) in the concentrations of TNFα in serum of diabetic animals compared to the control group (Table 6). In our opinion, the accumulation of proinflammatory cytokines detected in serum of diabetic animals displayed the development of systemic inflammation with insulin resistance to be the main attribute of T2DM. As confirmation to the previous statement, our data illustrated a significant correlation between serum levels of TNFα and the results of the OGTT (r^2^=0.88, p <0.01) in a T2DM group.

**Table 6.** The effect of betulin on serum TNFα levels in rats with T2DM.

| Groups | TNFα,<br>pg/ml |
| --- | --- |
| Control | 7.60±0.51 |
| Betulin | 5.98±0.65 |
| T2DM | 16.77±1.11<sup>A</sup> |
| T2DM+Betulin 50 | 15.01±1.21<sup>A</sup> |
| T2DM+Betulin 100 | 11.35±1.05<sup>B</sup> |

A – p <0.05 compared with the control group, B – p <0.05 compared with the T2DM group

The administration of betulin to diabetic animals did not fully recover high levels of TNFα associated with diabetes-induced inflammation and insulin resistance to the values obtained for non-diabetic rats. The lower dose of betulin (50 mg/kg/day) appeared to be less effective to abolish the accumulation of proinflammatory cytokines in serum of treated animals compared to non-treated diabetics and control animals. Nonetheless, the higher dose of betulin (100 mg/kg/day) partially alleviated diabetic conditions (an increase in pro-inflammatory marker by 49.4 %, p=0.0989) compared to control rats, and resulted in TNFα significantly reduced when compared to non-treated diabetic controls (Table 6). It should be noted that that the administration of betulin to non-diabetic rats was accompanied by a slight decrease in TNFα levels, which was 21.3 % lower than in the control group (p <0.1).

## Conclusion

It was shown for the first time that long-term administration of betulin to the experimental animals prevented insulin resistance and restored glycemic control in a dose-dependent manner in T2DM related conditions. Abnormal lipids in blood serum and histological changes in the liver tissues of rats were also reduced toward normal values. We suggested that normalization of lipid metabolism and the ability of betulin to restrain systemic inflammation can be accounted for improved insulin resistance and it can provide a possible mechanism for the beneficial antidiabetic effects of betulin.

## Acknowledgements

The authors would like to express their gratitude to A. A. Astrowski, Dr. Med. Sci., Prof. (Institute of Biochemistry of Biologically Active Compounds of the National Academy of Sciences of Belarus), N. I. Prokopchik, M.D., PhD. (Grodno State Medical University), L. Chyczewski, Dr. Med. Sci., Prof. (University of Medical Sciences in Bialystok) for their valuable help in the histopathological studies. The authors thanks to S. Szycko, M.D., PhD. (Medical University of Gdansk) for providing some of the reagents used in this study.

